# FGFR/Heartless and Smog interact synergistically to negatively regulate Fog mediated GPCR signaling

**DOI:** 10.1101/2019.12.30.890871

**Authors:** Kumari Shweta, Anagha Basargekar, Anuradha Ratnaparkhi

## Abstract

G-protein coupled receptor (GPCR) signaling triggered by Folded gastrulation (Fog) is one of the pathways known to regulate glial organization and morphogenesis in the embryonic CNS in *Drosophila*. Fog is best known for its role in epithelial morphogenesis during gastrulation. Here, the signaling pathway includes GPCRs Mist and Smog and the G-Protein Concertina (Cta) which activate downstream effectors to bring about cytoskeletal changes essential for cell shape change

In this study, we identify molecular players that mediate and serve as important regulators of Fog signaling in the embryonic CNS. We find that while Cta is essential for Fog signaling neither receptors, Mist nor Smog mediates signaling in the CNS. On the contrary, we find that Smog functions as a negative regulator of the pathway. Surprisingly, *Heartless* which encodes a fibroblast growth factor receptor, also functions as a negative regulator of Fog signaling. Further, we find that both *heartless* and *smog* interact in a synergistic manner to regulate Fog signaling.

This study thus identifies novel regulators of Fog signaling that may play an important role in fine-tuning the pathway to control cell morphogenesis. It also suggests the likelihood of there being multiple receptors for Fog that mediate and regulate signaling in a context specific manner.

**Author Summary:** In *Drosophila*, Folded gastrulation (Fog) functions as ligand that signals via GPCRs to regulate cell shape during gastrulation -one of the earliest events in embryogenesis. Here, Fog signals via receptors Mist and Smog to activate the G-protein Concertina to elicit change in cell shape. In the embryonic central nervous system (CNS) this pathway regulates shape and organization of glia important for functions such as insulation of neurons and synapses.

The mechanism of Fog signal transduction in the CNS and its regulation is not well understood. We have sought to address these questions in our study. We find that Concertina is an essential factor for Fog signaling in the CNS but interestingly Mist is not. In contrast, Smog functions as a negative regulator such that loss of Smog enhances Fog signaling. A similar role is played by the receptor tyrosine kinase-Heartless. Interestingly, we find that Smog and Heartless interact as part of a common genetic network to regulate Fog signaling. Our results thus provide novel insights into the regulation of Fog signaling and shed light on how signaling can be fine-tuned in a context dependent manner to control cell shape change which plays a critical role during development and organ formation.

## Introduction

Cells undergo change in shape during development to facilitate processes like cell migration, tissue extension and tube formation essential for organ formation. In *Drosophila*, GPCR signaling triggered by the ligand Folded gastrulation (Fog), is amongst the first to be activated to bring about co-ordinated cell shape change essential for gastrulation (Costa et al., 1994).

The signaling pathway consists of Concertina- the G*α*12/13 subunit of the heterotrimeric G-protein (Cta; Parks and Wiechaus., 1991), which activates RhoGEF2 and Rho Kinase eventually leading to actin cytoskeletal changes that facilitate cell-shape change (Dawes-Hoang et al., 2005). GPCRs namely Mist and Smog, have been identified as mediators of Fog signaling. Mist was identified as a receptor through a cell-culture based screen for suppressors of the pathway. This gene is zygotically expressed in early blastoderm embryos in a pattern similar to Fog (Manning et al., 2013). The second receptor called Smog, is maternally expressed and mediates part of the Fog signal during gastrulation (Kerridge et al., 2016). Activation of Fog signaling by these receptors leads to apical constriction which is lost once invagination is complete presumably through downregulation of the pathway. The details of this process are still unclear. The cells then turn mesenchymal through activation of FGF signaling mediated by the receptor Heartless (Htl; Leptin.,1999).

We are interested in understanding the mechanism of Fog signaling and its regulation in the embryonic central nervous system (CNS), particularly glia. In the embryonic CNS, *fog* mRNA expression is enriched in a subset of glia particularly the longitudinal or interface glial (LG), and in the peripheral nervous system (PNS) namely in chordotonal organ (CO), scolopale and cap cells. Loss of *fog* in glia alters morphology leading to defects in ensheathment of the neuropil, while overexpression of *fog* leads to disorganization of the glial lattice (Ratnaparkhi and Zinn.,2007). Interestingly, *heartless* is also expressed in LG (Shishido et al., 1997) and is shown to regulate extension of glial processes into the neuropil in the embryonic as well as larval CNS. The requirement of this signaling pathway has also been demonstrated in glia associated with larval eye-disc for ensheathing photoreceptor axons, and in the adult olfactory lobe for compartmentalization of olfactory glomeruli (Franzdóttir et al., 2009; Stork et al., 2014; Wu et al., 2017)

The extent of molecular conservation between Fog signaling in the CNS and the ventral furrow has not been determined. Further, molecular mechanisms involved in the regulation Fog signaling are still poorly understood. Through a wing based genetic screen, we identified regulators of mitochondrial fusion and fission as downstream modulators of Fog signaling (Ratnaparkhi, A., 2013). Mutations in *rptp52F* (*ptp52F*) which encodes an orphan receptor tyrosine phosphatase (Schindelholz et al., 2001), leads to formation of an irregular ventral furrow similar to *fog* mutants and genetic epistasis places *ptp52F* downstream of *fog* (Ratnaparkhi and Zinn.,2007). The genetic interaction between *ptp52F* and *fog* suggests that pathways involving tyrosine phosphorylation, are likely to regulate Fog signaling. Supporting this, we identified elements of the Heartless signaling as being epistatic to Fog in our screen (data not shown). The overlap in expression and function of Fog and Htl signaling in LG and their role in regulating morphogenesis, prompted us to test if Htl might regulate Fog signaling.

Overexpression of Fog leads to ectopic axonal and glial midline crossing in the embryonic CNS. Using this as an assay, we show that Fog signaling in the CNS requires Concertina: loss of *concertina* completely suppresses midline crossing making it an essential component of the pathway in the CNS. In contrast, we find that Mist is dispensable. Unexpectedly however, Smog in this context, functions as a negative regulator such that downregulation of *smog* results in enhanced Fog signaling. Interestingly, Htl also functions as a negative regulator. Furthermore, *htl* and *smog* interact genetically in a synergistic manner indicating that the two genes function as part of common genetic pathway to regulate Fog signaling. Supporting this, we find that Smog staining, but not mRNA levels, is reduced in *htl* mutants pointing towards a regulatory mechanism that is post-transcriptional.

Our study thus identifies *htl* as a novel regulator of Fog mediated GPCR signaling that functions by regulating the levels of Smog. It also highlights the context dependent function of Smog that might serve to fine-tune signaling in a tissue specific manner.

## Results

### Concertina is essential for Fog signaling in the CNS

Expression of *fog* is enriched in a subset of LG glia in the embryonic CNS and downregulation of *fog* leads to change in glial size and morphology: the LG become smaller and more spherical in shape (Ratnaparkhi and Zinn., 2007; Compare Fig 1A and 1B). We knocked down *concertina* (*ctaRNAi*) in LG using RNAi to determine whether it exerts a similar effect on LG morphology. Similar to *fogRNAi*, LG in these embryos appeared small and more spherical (Fig. 1C). A measurement of the aspect ratio of these cells revealed a small but significant decrease of 13% compared with control (Fig. 1D).

**Figure 1.**
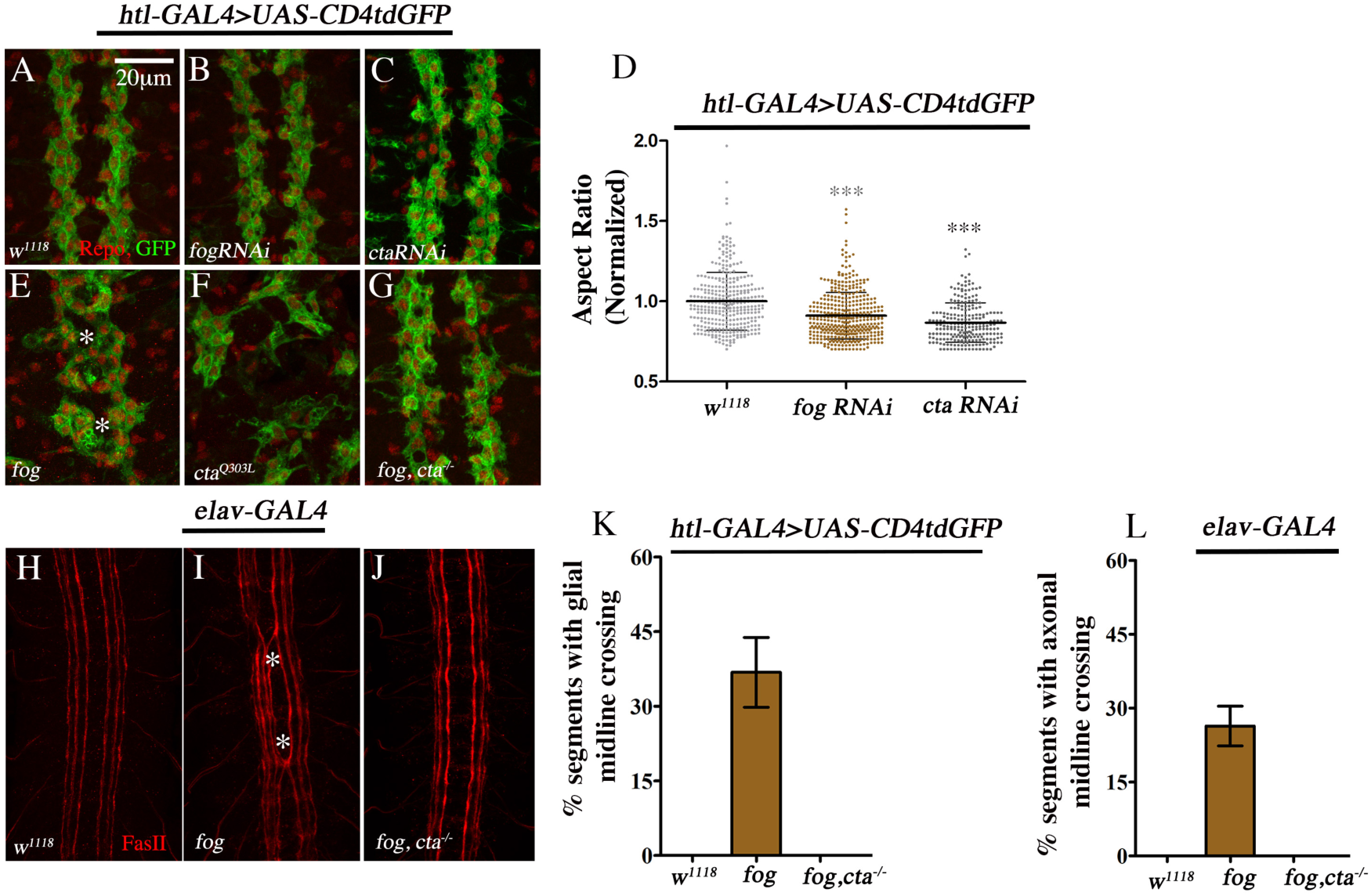
Fog regulates morphology and organization of longitudinal glia. (LG). (A-G) Longitudinal glia in the embryonic CNS stained with GFP (green) and Repo (red) which mark the glial cell membrane and nuclei respectively. (A) CNS of control embryo showing LG organized on either side of the midline. The cells appear ovoid. (B-C) Knockdown of *fog* (B) and *cta* (C) using RNAi alters cell shape causing cells to appear small and spherical. (D) Quantification of glial aspect ratio (Control: 1.0±0.18 (n=337, N=17) versus *UAS-fogRNAi*: 0.91±0.15 (n=408, N=20) versus *UAS-ctaRNAi* 0.87±0.12 (n=240, N=12)). (E) Overexpression of *fog* leads to glial disorganization. LG form clusters, appear stretched and can be seen extending processes across the midline (asterisk). (F) Overexpression of constitutively active *cta* leads to a complete disorganization of the glial lattice. (G) Overexpression of *fog* in a *cta* mutant background leads to a complete suppression of glial midline crossing. (H-J) CNS stained with anti-FasII. Control embryos (H) at late stage16 show 3 axon fascicles present on either side of the midline. Overexpression of Fog leads to ectopic axonal midline crossing (I, asterisk) which is suppressed upon loss of *cta* (J). (K) GMC in *htl*-GAL4>*UAS-fog* is 36.81±7.02%. Loss of *cta* suppresses GMC completely. (L) AMC in *elav*-GAL4>*UAS-fog* embryos is 26.37±4.02% which is completely suppressed in a *cta* mutant. Error bar in (D) represents SD; in K and L the error bar represents SEM. *** indicates P<0.0001.

Overexpression of *fog* in glia, leads to disorganization of the glial lattice and embryonic lethality (Ratnaparkhi and Zinn., 2007). In terms of morphology, LG appear stretched and more tightly packed. A closer look at the disorganization revealed that in some segments, LG were present either at the midline or closer to the midline with processes extending across (Fig. 1E, asterisk). We refer to this phenotype as ectopic ‘glial midline-crossing’ or GMC for short. Interestingly, overexpression of *concertina* did not significantly alter glial organization; however, expression of a constitutively active Concertina (*UAS-cta^Q303L^*/ *UAS-cta^CA^*) led to severe defects: LG were scattered and appeared stretched. The CNS of these embryos failed to condense (Fig. 1F).

We carried out genetic epistasis experiments to determine whether *concertina* is essential for Fog signaling using the GMC as an assay. Indeed, overexpression of *fog* in a *concertina* mutant led not only to a completely suppression of GMC, but also rescued the lethality caused by *fog* overexpression. Compared with 37% GMC in *UAS-fog* embryos, we did not observe any crossovers in the absence of *concertina*. (Fig. 1G, K).

*Fog* is also expressed in neurons and seen to regulate axon guidance (Ratnaparkhi and Zinn., 2007). Pan-neuronal overexpression of *fog* in neurons leads to ectopic axonal midline crossing (AMC). As with GMC, loss of *concertina* led to a complete suppression of AMC (Fig.1J, L) compared with the 26% seen in *elav-*GAL4>*UAS-fog* embryos (Fig. 1I; Ratnaparkhi and Zinn., 2007). Together these results indicate that Concertina is essential for transducing the Fog signal in the embryonic CNS.

### Smog is expressed in the embryonic CNS

Fog is known to bind to two G-protein coupled receptors (GPCRs) encoded by genes *mist* and *smog* (Manning et al., 2014; Kerridge et al., 2016). ModENCODE data shows that the latter is expressed at high levels and almost exclusively in the CNS (www.flybase.org). Based on this, we sought to first test if Smog functions as a receptor for Fog in the embryonic CNS. As a first step, we examined the expression pattern of Smog in the CNS using an antibody raised against the N-terminal region of the protein.

*Smog* is expressed maternally and is present ubiquitiously in blastoderm embryos (Kerridge et al., 2016). Consistent with this, we observed a uniform and punctate Smog staining in early blastoderm embryos (Fig. 2A). Later, during gastrulation, a more localized expression was detected in invaginating cells of the ventral furrow (Fig. 2B). In the nervous system, Smog expression appeared remarkably similar to Fog with staining seen in the CNS (Fig. 2C) and peripheral chordotonal organs (CHO; Fig. 2E and 2E’). The intensity of staining was reduced upon knock-down of *smog* using RNAi (Fig. 2D, 2F and 2F’). We quantified the decrease in staining by measuring fluorescence intensity at the CHO. Compared to control, embryos expressing *smogRNAi* showed a 40% decrease in staining intensity (Fig. 2G) while in *smog* mutants (*smog^KO^;* Kerridge et al., 2016), the intensity was reduced by 60% (Fig. 2G). Smog staining was also reduced in *Dmon1^Δ129^*/*Df(2L)9062* embryos (Compare Fig. S1A and S1B) in which *Dmon1^Δ129^* represents a small deletion in the region spanning the 3’ region of *mon1* (CG11926; Deivasigamani et al., 2015) and 5’ upstream sequence of *smog*. Immunohistochemistry with the pre-immune serum did not exhibit any specific staining in the CNS or the PNS (Fig. S1C and S1D).

**Figure 2.**
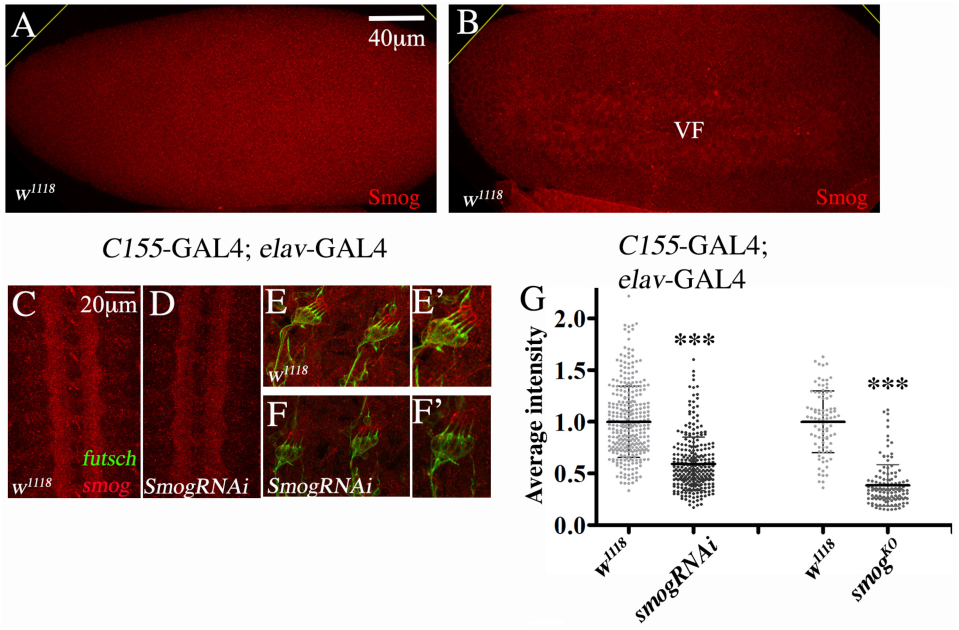
Smog is expressed in the embryonic CNS and PNS. (A-B) Wild-type embryo at the blastoderm (A) and gastrulation (B) stages. Note the expression of Smog (red) in the region of the ventral furrow (VF) in B. (C-D) CNS of control (*C155*-GAL4/+; *elav*-GAL4/+; C) and *UAS-smogRNAi* embryo (*C155-*GAL4*; elav-*GAL4*>UAS-smogRNAi*; D). Note the reduced staining in (D). (E-F) Smog staining in chordotonal organ (CHO) of control (E&E’) and *UAS-smogRNAi* embryos (F&F’). Note the reduced staining in (F). (G) Quantification of the fluorescence intensity in the CHO of *smogRNAi and smog^KO^* mutants. A 40% and 60% decrease in staining is seen in RNAi and mutant embryos respectively (*C155*-GAL4/+; *elav*-GAL4/+: 1±0.02, n=315 versus *C155-*GAL4*; elav-*GAL4*>UAS-smogRNAi*: 0.59±0.01, n=274; *w^1118^:* 1.0±0.30 n=92 versus *smog^KO^*: 0.38±0.20 n=126). n= number of neurons. *** indicates p<0.0001

### Smog functions as a negative regulator of Fog signaling in the CNS

To determine whether Smog functions as the Fog receptor in the CNS, we checked if loss of *smog* suppressed AMC caused by *fog* overexpression. Surprisingly, knockdown of *smog* using RNAi led to a significant increase in midline-crossing. Compared with *elav*-GAL4>*UAS-fog* (Fig. 3A and 3G), embryos co-expressing *UAS-smogRNAi* showed a 70% increase in AMC (Fig. 3B & 3G). This was completely suppressed in the absence of *cta* (Fig. 3C & 3G) indicating that the enhanced phenotype is indeed due to upregulation of Fog signaling.

**Figure 3.**
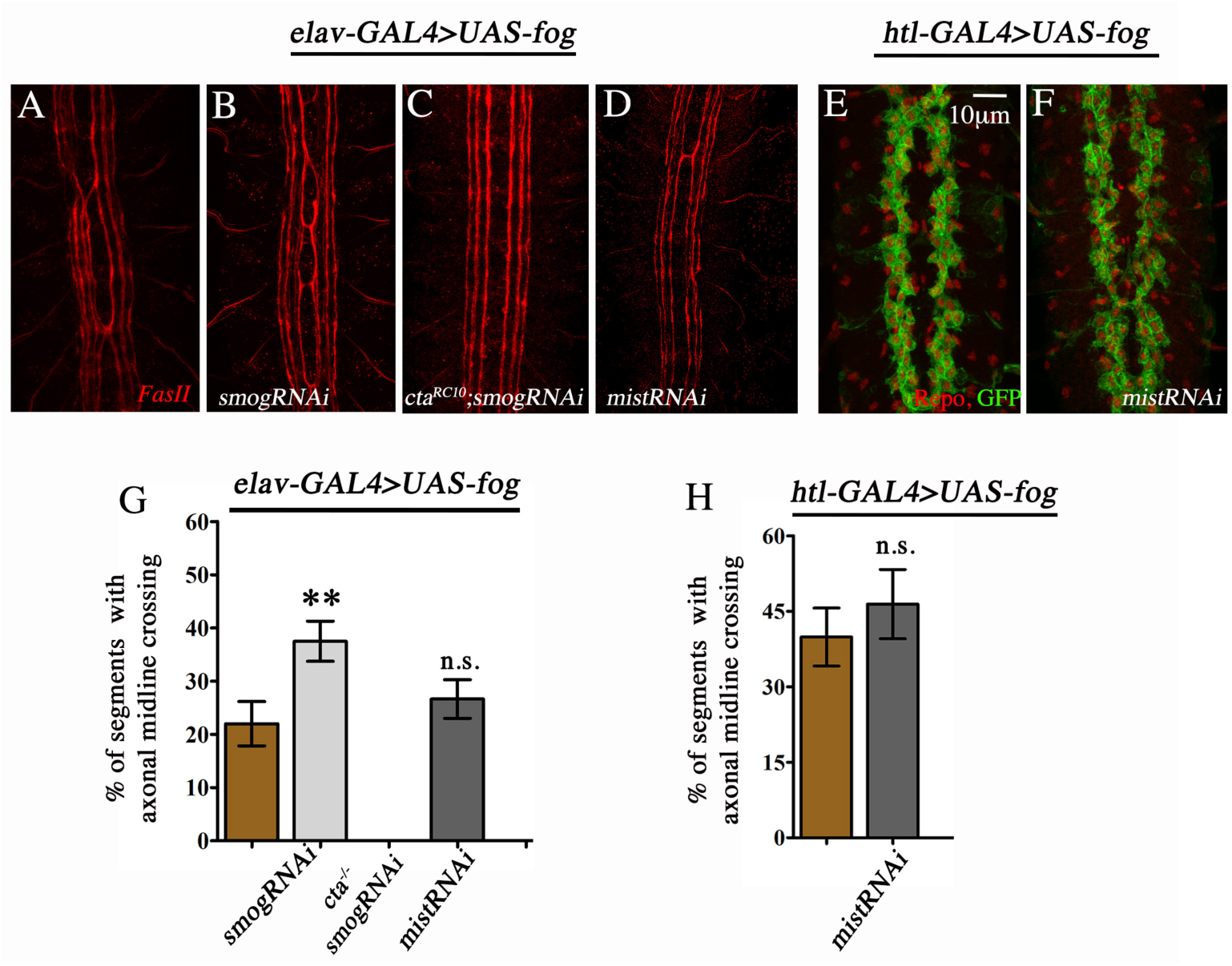
Smog is a negative regulator of Fog signaling in the CNS. (A-D) Embryonic CNS stained with anti-FasII (red). (A) AMC in *elav*-GAL4>*UAS-fog* embryo. (B) Expression of *smogRNAi* enhances AMC. (C) Loss of *cta* completely suppresses AMC. (D) Downregulation of *mist* does not enhance AMC. (E-F) Embryos expressing *UAS-fog* (E) and *UAS-fog, UAS-mistRNAi* in glia (F) show comparable GMC. (G) Percentage AMC: *elav*-GAL4>*UAS-fog*: (22.01±4.16, n=259); *elav*-GAL4>*UAS-fog, UAS-smogRNAi* (37.05±3.78, n=224); *elav*-GAL4>*UAS-fog*, *cta^RC10^* (0, n= 259); *elav*-GAL4>*UAS-fog, UAS-mistRNAi* (26.64±3.64, n= 259). (H) Percentage GMC: *htl*-GAL4>*UAS-fog* (38.90±5.75; n=198) and *htl*-GAL4>*UAS-fog, UAS-mist RNAi* (46.43±6.89; n=168). n= number of segments.

Given the unexpected result with *smog*, we sought to test whether Mist functions as the Fog receptor. However, knock-down of *mist* did not alter AMC nor GMC (Compare Fig. 3A and 3D; 3E and 3F respectively). The penetrance of these phenotypes in embryos expressing *UAS-mistRNAi* was comparable to their respective controls (Fig. 3G and 3H). The above results thus reinforce the role of Concertina (G*α*12/13) as an essential component of the Fog pathway in the CNS and indicate that while neither Mist nor Smog is involved in transducing the Fog signal, the latter functions as a negative regulator by restricting signaling.

### *Heartless* negatively regulates Fog signaling

Loss of *fog* leads to defects in the glial organization and morphology; the axonal tracts in the CNS show breaks (Ratnaparkhi and Zinn., 2007) - a phenotype also observed in other mutants with defects in glial organization such as *deadringer* (*dri*), *fear-of-intimacy* (*foi*) and *heartless* (*htl*; Shandala et al., 2003; Pielage et al., 2004; Shishido et al., 1997). For reasons stated earlier and because of its role in glia, we chose to test whether Htl regulates Fog signaling. As a first step, we examined LG organization in *htl* mutants to determine the extent of similarity, if any, to *fog* loss-of-function and gain-of-function embryos. In *htl^AB42^* - a null mutant of the gene (Gisselbrecht et al., 1996) LG appeared small and more clustered. The glia seemed to adhere more closely in a manner that was more similar to *fog* overexpression than *fogRNAi* or *cta* mutants. Occasional breaks in the glial tracts were also observed (Compare Fig. S2A and S2B). The tendency to cluster was also observed upon knock-down of *htl* using RNAi (Fig. S2C). In terms of organization, the LG were mildly disorganized. A detailed comparison of LG size in *htl^AB42^* mutants, and embryos expressing *UAS-htlRNAi* revealed a very modest (5%) decrease. Interestingly, neither overexpression of constitutively active *htl* (*λ*htl) nor the ligand *thisbe*, altered organization, size or aspect ratio of the LG. Hyperactivation of the pathway was confirmed by the robust increase in glial number seen upon driving expression with *repo*-GAL4.

Using GMC and AMC as an assay, we examined whether Fog and Htl signaling pathways interact genetically. Interestingly, co-expression of UAS-*htlRNAi* with *UAS-fog* in glia led to a strong increase in GMC: the phenotype was observed in 51% of the segments compared to 38% in *UAS-fog* embryos (Fig. 4B and 4D). In a *htl^AB42^* mutant, 63% of the segments showed GMC (Fig. 4C and 4C’ and 4D). Further, LG in these embryos seemed tightly packed (Fig. 4B and 4C) and, in more severe cases, the clusters localized at the midline (Fig. 4C’, arrow). Enhanced GMC was also seen in *htl^AB42/YY262^* and *htl^YY262^* mutants (65% and 67% respectively; Fig. 4D). Consistent with these results, compared with controls (*htl-*GAL4, *UAS-Cd4tdGFP*>*UAS-fog*), the number of embryos with >3 GMC was considerably more in the mutant pool (Fig. 4E). In *htl* mutants alone, GMC was observed at a low frequency of 5%. This, together with the above results suggests that loss of *htl* enhances the phenotype of *UAS-fog* and thus functions as a negative regulator of the Fog signaling.

**Figure 4.**
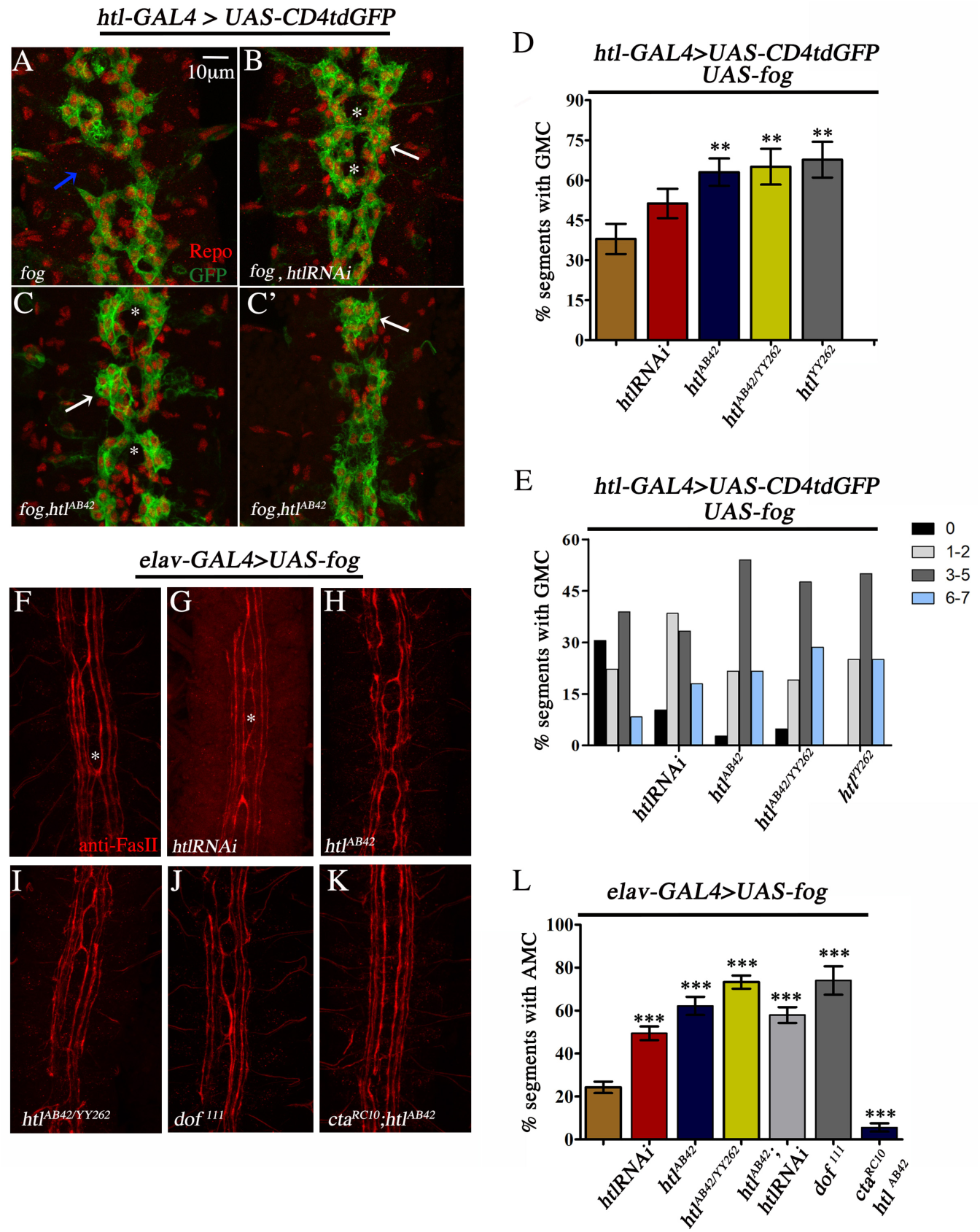
Loss of *htl* enhances the AMC and GMC phenotypes caused by Fog overexpression. (A) GMC and LG clustering in *htl-GAL4>UAS-fog* embryo. Occasional gaps seen in the LG scaffold is marked (blue arrow). (B) Knock-down of *htl* in glia using RNAi, leads to increased clustering and enhanced GMC. (C and C’) GMC is enhanced in a *htl^AB42^* mutants. In more severe phenotypes, LG cluster at the midline (C’, arrow). (D) Quantification of GMC in *UAS-fog*: (37.96±5.66, n=216); *htlRNAi*: (51.28±5.52, n=234); *htl^AB42^*: (63.06±5.13, n=222); *htl^AB42/YY262^*:(65.08±6.70, n=126); *htl^YY262^*: (67.71±6.71, n=96). (E) Percentage distribution of GMC in *UAS-fog* (0=30.56; 1-2=22.22; 3-5=38.89; 6-7=8.33); *htlRNAi*: (0=10.26; 1-2=38.46; 3-5=33.33; 6-7=17.95); *htl^AB42^*: (0=2.70; 1-2=21.62; 3-5=54.05; 6-7=21.62); *htl^AB42/YY262^*: (0=4.76; 1-2=19.05; 3-5=47.62; 6-7=28.57); *htl^YY262^*: (0=0; 1-2=25; 3-5=50; 6-7=25). (F-K) CNS of embryos stained with anti-FasII. (F) AMC in *elav*-GAL4>*UAS-fog* is marked by an asterisk. Note the increase in AMC upon expression of *htlRNAi* (G) and in *htl^AB42^* (H), *htl^AB42/YY262^* (I) and *dof^111^* mutants (J). AMC is suppressed in a *cta^RC10^*; *htl^AB42^* double mutant (K). (L) Percentage AMC in *UAS-fog*: (24.24±2.67, n=231); *htlRNAi*: (49.41±3.16, n=427); *htl^AB42^*: (62.18±4.23, n=119); *htl^AB42/YY262^*:(73.26±3.07, n=273); *htl^AB42/htlRNAi^*: (57.93±3.69, n=252); *dof^111^*: (74±6.62, n=77); *cta^RC10^*; *htl^AB42^*: (5.58±1.92, n=287).

### Downstream-of-FGF (Dof) is a negative regulator of Fog signaling

*Fog* and *htl* also function in neurons: both genes are known to play a role in axon guidance (Forni et al., 2004; Ratnaparkhi and Zinn., 2007). Based on this, we sought to test whether they interact in similar manner in the neuronal context as well. Similar to the results with GMC, co-expression of *UAS-htlRNAi* with *UAS-fog* led to a near two-fold increase (49%) in AMC. (Compare Fig. 4F & 4G; Fig. 4L). A significant increase in AMC was also observed in *htl^AB42^* and *htl^AB42/YY262^* mutants resulting in 62 % and 73% midline crossing respectively (Fig. 4H-I and 4L).

*Stumps*/*downstream-of-fgf (dof*) is a specific effector of FGFR signaling in *Drosophila* (Vincent et al., 1998). Consistent with the above results, loss of *dof^111^* led to an increase in AMC comparable to *htl* mutants: 74% of the segments showed midline-crossing (Fig. 4J & 4L). This indicates that the signaling pathway mediated by Htl, and not just the receptor, regulates Fog signaling.

We confirmed that the observed increase in ‘midline-crossing’ was due to upregulation of Fog signaling, by expressing *UAS*-*fog* in a double mutant lacking both *cta* and *htl* (*cta^RC10^*; *htl^AB42^*). Indeed, midline-crossing was strongly suppressed in these embryos (5.6 %; Fig. 4K and 4L) thus emphasizing the central role of Cta in Fog signaling.

Based on the above results, we wondered whether expression of constitutively active *htl* (*UAS-λhtl*) might suppress ‘midline-crossing’. Surprisingly, we observed that the phenotype is enhanced (Fig. S3B & S3E). A similar result was obtained upon co-expression with *UAS-thisbe::HA (ths)* and *UAS-htl*, although the extent of increase in AMC in case of the latter seemed less compared with the others (Fig. S3C & S3E). Notably, individual expression of neither *UAS-λhtl* nor *UAS-thisbe::HA or UAS-htl* led to AMC. While one cannot rule out promiscuous signaling as a reason for the enhancement, this suggests Htl needs to signal at a certain optimal level in order to modulate the Fog pathway; too much or too little signaling leads to upregulation of the Fog pathway. Together, the results described above indicate that Htl signaling functions to restrict Fog activity in the embryonic CNS.

### The interaction between *fog* and *htl* is cell autonomous

Being a secreted ligand, overexpression of Fog in glia also affects neurons resulting in ectopic axonal midline crossing albeit at a lower frequency (Ratnaparkhi and Zinn., 2007). To test if *htl* can influence Fog signaling in a non-cell autonomous manner, we co-expressed *UAS-fog* with *UAS-htlRNAi* in glia and examined its effect on AMC. In *htl*-GAL4>*UAS*-*fog* embryos, the frequency of AMC is approximately 8 % (Fig. 5B & 5G). This is completely suppressed upon loss of *cta,* which further confirms that it is essential for signaling (Fig. 5C). Interestingly, glial expression of *UAS-fog* and *UAS-htlRNAi* did not enhance AMC (12%; Fig. 5D & 5G); the distribution of AMC in these embryos was similar to the controls (*htl*-GAL4>*UAS*-*fog*; Fig. 5H). In contrast, glial overexpression of *UAS-fog* in *htl^AB42^* and *htl^AB42/YY262^* led to enhanced AMC (42 % and 52% respectively; Fig. 5E-G) with a sharp increase in the number of embryos with more than 3AMC per embryo (Fig. 5H). Thus, co-expression of *UAS-fog* with *UAS-htlRNAi* in glia does not enhance AMC, whereas a significant increase is observed in a *htl* mutant. This latter result thus indicates that the increase in AMC must arise due to interaction between *fog* and *htl* in neurons. Based on this, we conclude that Htl functions in a cell autonomous manner to regulate Fog signaling.

**Figure 5.**
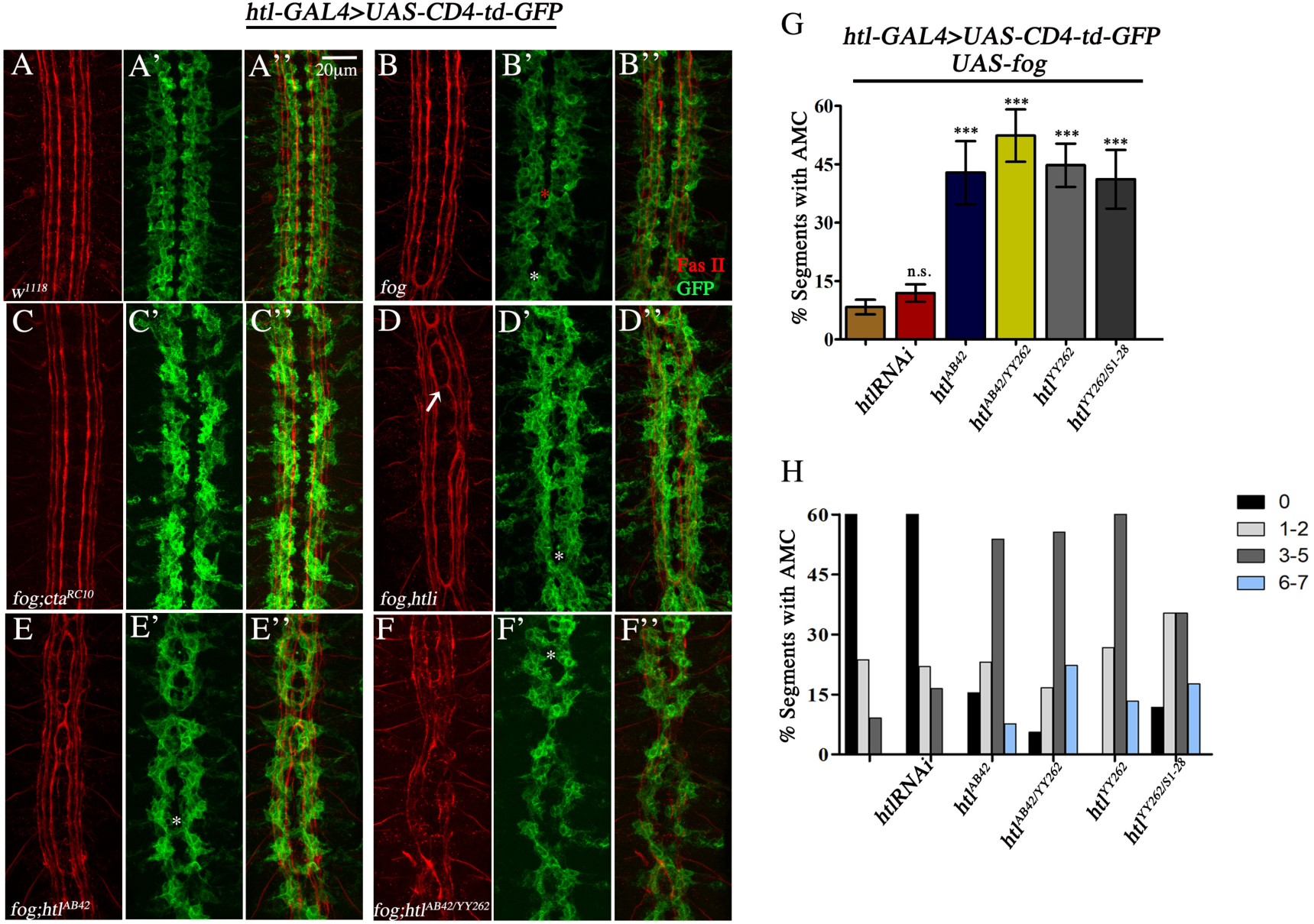
Interaction between *fog* and *htl* is cell autonomous. (A-A’’) Control embryo (*htl-*GAL4*, UAS-CD4tdGFP/+*) stained with anti-FasII and anti-GFP. Note the organization of glia and axons. (B-B’’) Overexpression of *fog* using *htl*-GAL4 leads to glial and axonal midline crossing which overlap with each other (white asterisk). Occasionally only GMC is observed (red asterisk). (C-C’’) Loss of *cta* completely suppresses GMC and AMC (D-D’’) Co-expression of *UAS-htlRNAi* with *UAS*-*fog* in glia does not lead to any significant enhancement in AMC. (E-F) Glial overexpression of *fog* in *htl^AB42^* and *htl^AB42/YY262^* mutants enhances AMC. (G) Quantification of AMC: *htl-GAL4>UAS-fog* (control, 8.31±1.88, n=385); *htl-GAL4>UAS-fog; UAS-htlRNAi* (11.94±2.23, n=511); *htl^AB42^* (42.86±8.09%, n=91); *htl^AB42/YY262^* (52.38±6.73, n=126); *htl^YY262^* (44.76±5.53, n=105); *htl^YY262/S1-28^* (41.18±7.54, n=119). (H) Percentage distribution of AMC: *UAS-fog* (0=67.27; 1-2=23.64; 3-5=9.09; 6-7=0); *UAS-htlRNAi*: (0=61.64; 1-2=21.92; 3-5=16.44; 6-7=0); *htl^AB42^*: (0=15.38; 1-2=23.08; 3-5=53.85; 6-7=7.69); *htl^AB42/YY262^*: (0=5.56; 1-2=16.67; 3-5=55.56; 6-7=22.22); *htl^YY262^*: (0=0; 1-2=26.61; 3-5=60; 6-7=13.3); htl^S1-28/YY262^ (0=11.76; 1-2=35.29; 3-5= 35.29; 6-7=17.65). n= number of segments. ***P<.0001, P≥0.05=n.s.

### *Htl* and *smog* interact in a synergistic manner to regulate Fog signaling

How does Htl regulate Fog signaling? The above results show that loss of *cta* can suppress the enhanced AMC and GMC phenotype due to loss of *htl*. This suggests that the regulation by Htl is likely to be at the level of the Fog receptor (Fig.6A). Given that both Htl and Smog have a similar effect on Fog signaling, we checked if the two genes interact to regulate the pathway. To test this, we expressed *UAS*-*fog* in neurons using *C155*-GAL4, which produces a weak AMC phenotype in about 6% of the segments (Fig 6B and 6E). Knockdown of *smog* using RNAi led to a small but significant enhancement in AMC (13%; Fig. 6E). A stronger enhancement of the phenotype was observed upon co-expression with *smogRNAi* in a *smog^KO/+^* mutant (43%; Fig. 6C and 6E).

**Figure 6.**
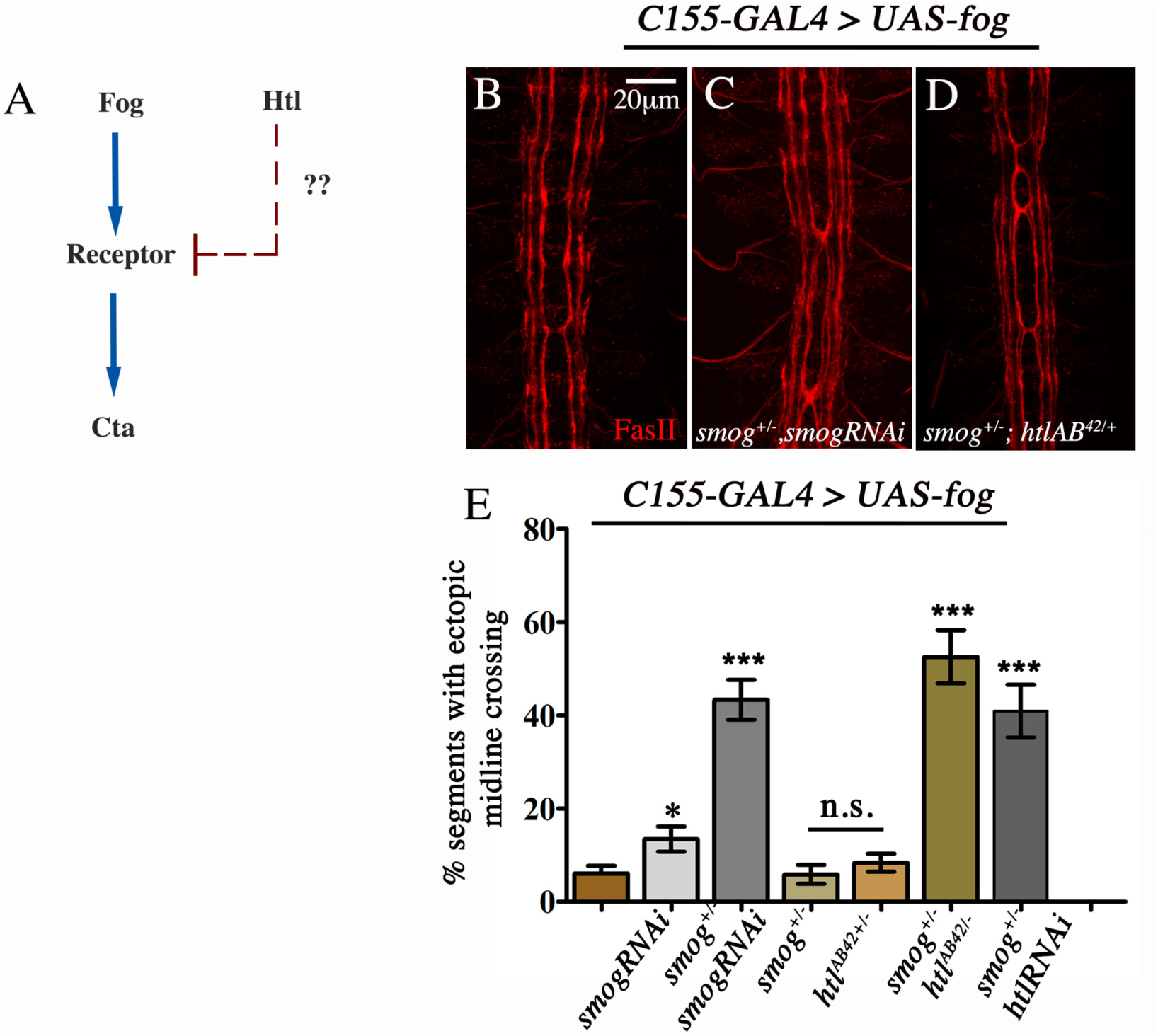
*Smog* and *htl* interact in a synergistic manner to regulate Fog signaling. (A) The interaction between Fog and Htl signaling pathways predicts that the regulation is likely to be at the level of the receptor. (B-D) Embryonic CNS stained with anti-FasII (red). AMC in *C155*-GAL4>*UAS-fog* embryo (B) and *C155*-GAL4>*UAS-fog,smog^KO/+^,UAS-smogRNAi* embryo (C). Note the increase in AMC. (D) *C155*-GAL4>*UAS-fog, smog^KO/+^, htl^AB42/+^*. Loss of one copy each of *smog* and *htl* enhances AMC. (E) Percentage AMC: *UAS-fog* (Control): 6.06±1.65 % (n=231)*; UAS-smogRNAi*: 13.45±2.69% (n=238); *smog^KO/+^;UAS-smogRNAi*: 43.35±4.28% (n=203); *smog^KO/+^*: 5.8±2.03% (n=154); *htl^AB42^*^/+^: 8.36±1.93% (n=287); *smog^KO^*^/+^*;htl^AB42^*^/+^: 52.57±5.7% (n=175); *UAS-fog*, *smog^KO^*^/+^*;UAS-htlRNAi*: 40.91±5.66% (n=154). n= number of segments. *** indicates P<0.0001, * indicates P<0.05; P≥0.05=n.s.

To test if *htl* and *smog* interact in a synergistic manner, we scored for AMC in embryos that were heterozygous for both *smog* and *htl* (*smog^KO^/+; htl^AB42^/+*). Interestingly, while neither loss of one copy of *smog^KO^* or *htl^AB42^* had any significant effect on AMC (6%, and 8% respectively; Fig. 6E), a dramatic increase was observed in *smog^KO^/+; htl^AB42^/+* embryos with 53% of the segments exhibiting AMC-a frequency comparable to that observed in *elav*-GAL4>*UAS-fog*; *htl^AB42^* embryos (Fig. 6D & 6E; Compare Fig. 6E and Fig. 4L). A similar increase in AMC was also seen upon expression of *UAS-htlRNAi* in a *smog^KO/+^* background (Fig. 6E) indicating that *htl* and *smog* interact synergistically to regulate Fog signaling.

### Loss of *htl* downregulates *smog* expression

The above results suggest that *htl* might function by regulating Smog levels. Supporting this possibility, we observed reduced Smog staining in *htl* mutants. (Compare Fig. 7A with 7B; Fig. 7C with 7D-F). A measurement of the fluorescence intensity at the CHO revealed a 35% decrease in both, *htl^AB42^* and *htl^YY262^* embryos (Fig. 7I). A comparable decrease in staining was observed in embryos expressing *UAS-htlRNAi* (Compare Fig. 7G & H; 7I). Further supporting this, we observed reduced Smog staining in *dof* mutants (28%; Fig. S4). Interestingly, we did not observe any significant increase in Smog staining in embryos expressing *UAS-λhtl* or *UAS-thisbe* (Fig. S5) indicating that the control of Smog levels is unlikely to be due to a transcriptional regulation by *htl*. To test this, we checked *smog* mRNA levels in wildtype and *htl^AB42^* mutant embryos using quantitative RT-PCR (qPCR).

**Figure 7.**
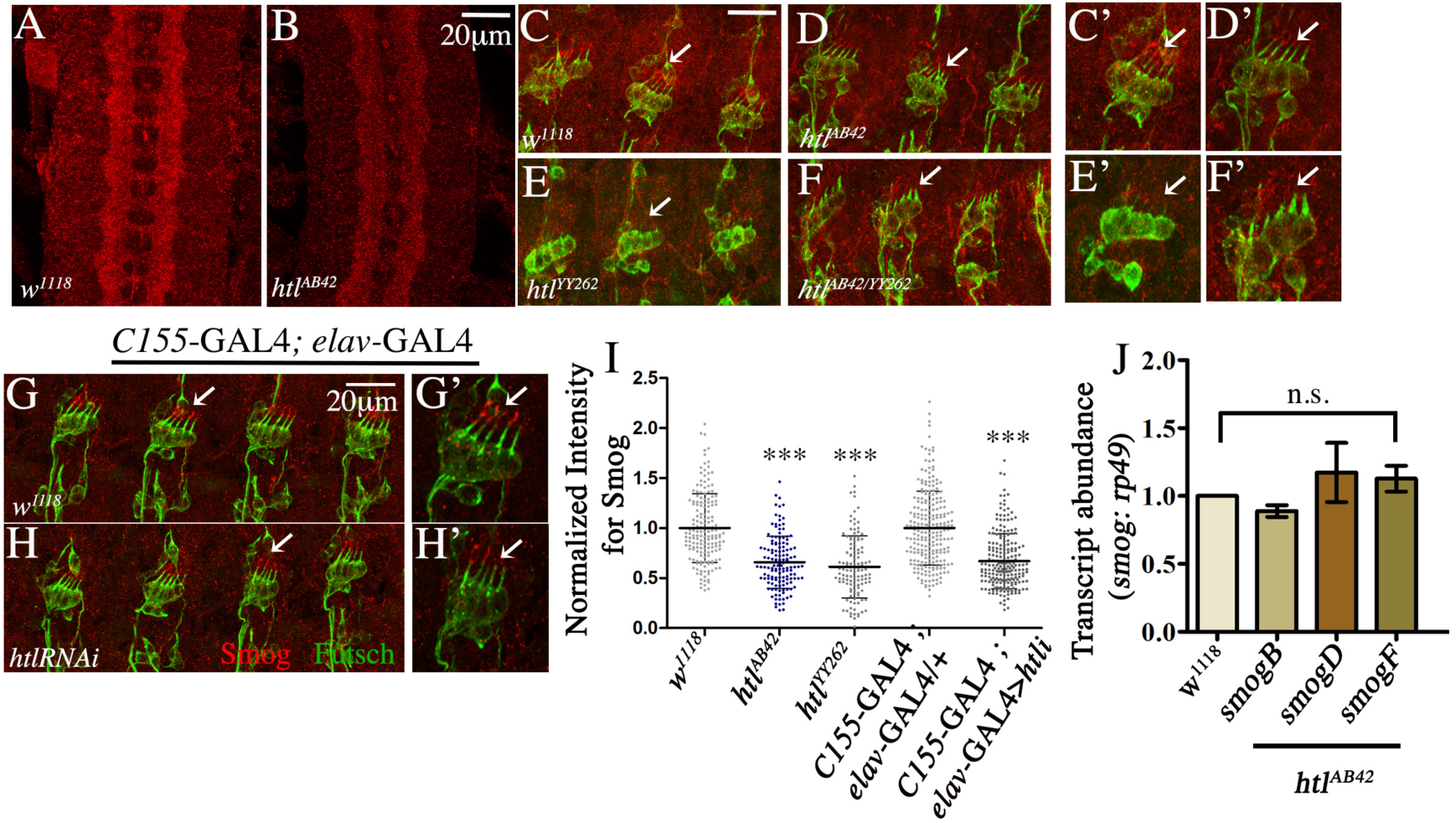
Smog staining is reduced in *htl* mutants. (A-B) CNS of wildtype (A) and *htl^AB42^* (B) embryos. Note reduced staining in B. (C-F) Smog expression (red) in CHO (anti-futsch, green) of *w^1118^* (C, C’), *htl^AB42^* (D, D’), *htl^YY262^* (E, E’) and *htl^AB42/YY262^*(F, F’). Smog levels are reduced in *htl* mutants. (G-H) Smog expression in CHO of *C155*-GAL4/+; *elav*-GAL4/+ (Control, G) and *C155*-GAL4/+; *elav*-GAL4/+>*UAS-htlRNAi,* embryos (H). Reduced staining is seen in the latter. (I) Normalized mean gray value for Smog intensity at the CHO. A significant decrease is seen in *UAS*-*htl* RNAi and *htl* mutants: (*w^1118^*: 1±0.34, N=13, n=171; *htl^AB42^*: 0.66±0.26, N=12, n=136; *htl^YY262^*: 0.60±0.31, N=10, n=118; *C155*-GAL4/+; *elav*-GAL4/+: 1±0.39, N=18, n=240; *C155*-GAL4; *elav*-GAL4>*UAS*-*htlRNAi*: 0.69±0.28, N=17, n=214). (J) Quantification of *smogB*, *smogD* and *smogF* transcripts in *htl^AB42^* mutants using qPCR: *w^1118^*: 1.0; *smogB*: 0.89±0.04; *smogD*:1.17±0.22; *smogF* 1.13±0.10. The values shown in (I) are mean±SD; values in (J) are mean±SEM. N= number of embryos and n= number of neurons. *** indicates P<0.0001.

**Figure 8.**
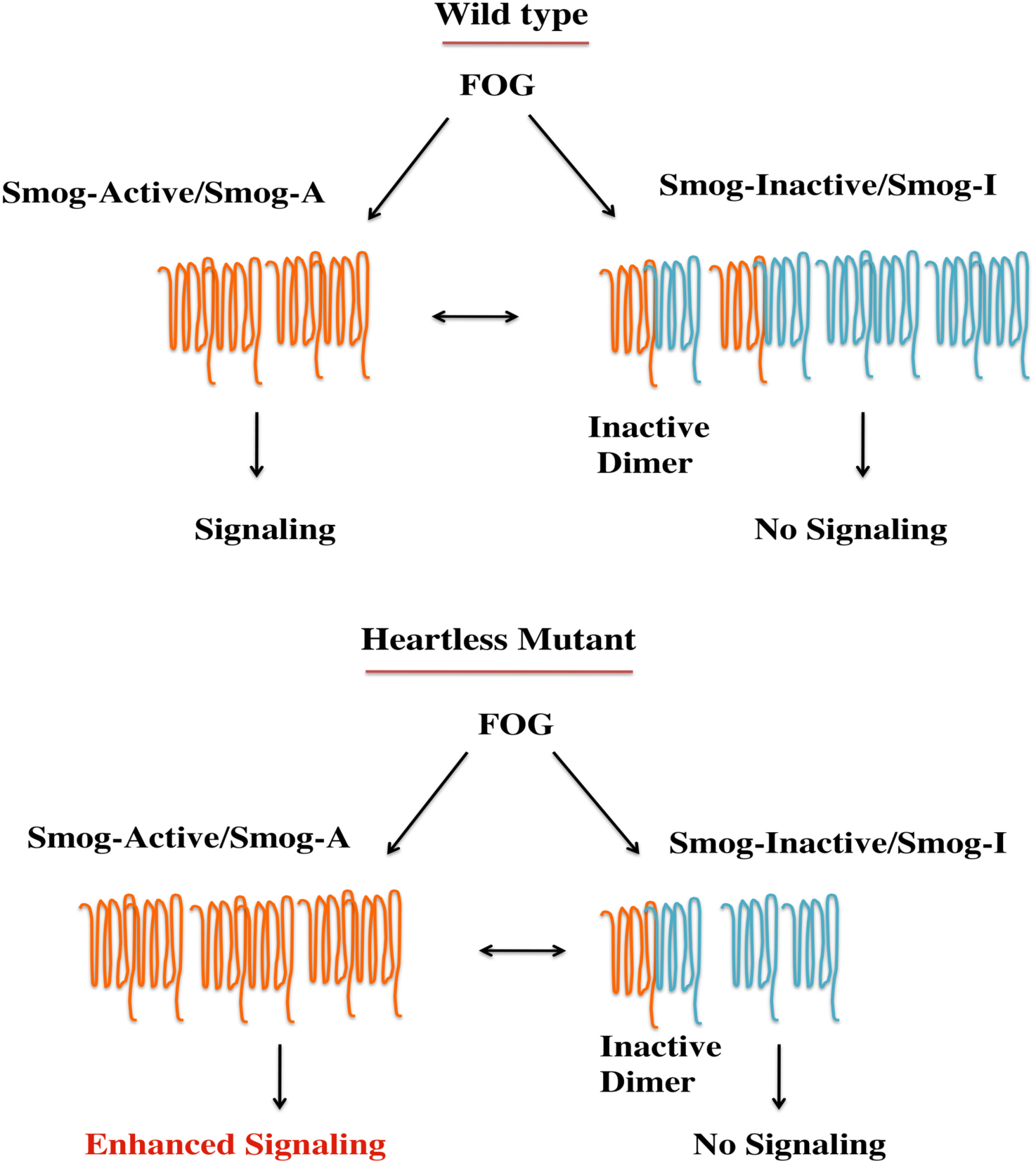
Proposed model to explain regulation of Fog signaling by Htl. We hypothesize that an inactive variant of Smog referred to as Smog-I is expressed in the CNS. Smog-I binds Fog but does not transduce the signal. Dimers with Smog-I are thus inactive. Htl signaling helps maintain a dynamic balance between Smog-I and the ‘active’ Smog-A. Absence or downregulation of Htl signaling destabilizes Smog-I not only making more ligand available for signaling but also shifting the receptor balance towards more Smog-A which leads to enhanced signaling.

*Smog* encodes multiple splice variants which, at the protein level, differ from each other at the C-terminus. Of these, the mRNA for *smogF* is the longest while *smogB* is the smallest with a short C-terminal domain that is unique compared to other isoforms. Using primers specific to *smog B*, *D* and *F* variants, we measured both, the relative abundance of these variants in wildtype embryos and the change in expression, if any, in *htl* mutants. The results from the qPCR revealed that *smog* splice variants are differentially expressed in the embryo with *smogF* being the most abundant of the three (Fig. S6). They also showed that mRNA levels of these transcripts are not significantly altered in *htl^AB42^* mutants (Fig.7J). As a control, we carried out qPCR to compare the transcript level of *htl* in the mRNA isolated from wildtype and *htl^AB42^* mutants and found a 70% reduction in *htl* mutants (Fig.S6). Thus, *htl* does not appear to regulate *smog* transcription-a result that also explains the absence of enhanced Smog staining in embryos expressing *UAS-λhtl*. This also suggests the involvement of a post-transcriptional mechanism in the regulation of Smog levels. These results, together with the genetic data discussed above, show that signaling by Htl helps restrict the Fog pathway by regulating Smog which functions as a negative regulator of Fog signaling in the CNS.

## Discussion

Fog activates one the earliest zygotic signaling pathways during development to co-ordinate cell shape for invagination of the presumptive mesoderm (Costa et al., 1994). It is also known to play a role in the development of the salivary glands, the adult wing, axon guidance and glial morphogenesis in the embryonic CNS (Lammel and Saumweber., 2000; Nikolaidou and Barrett., 2004; Ratnaparkhi and Zinn., 2007). The extent to which this signaling pathway is conserved in these different developmental contexts and its mode of regulation needs to be understood. We have addressed some of these questions in the context of the embryonic CNS.

It is established that Concertina is essential for Fog signaling during gastrulation (Morize et al., 1998). Our study shows that this holds good in the context of the CNS as well, given that both AMC and GMC due to Fog overexpression is completely suppressed in a *concertina* mutant (Fig. 1). In addition, the lethality caused by Fog overexpression in glia is also suppressed.

A key finding here is the interaction between *fog* and *htl* which signal to regulate morphogenesis, first during gastrulation (Leptin.,1999) and later, in the LG (Shishido et al.,1997; Stork et al., 2014). The role of Htl signaling appears to be context dependent. In the embryo, consistent with the findings of Stork and colleagues (Stork et al., 2014), we do not find any change in the number of Repo and Prospero positive LG in *htl* mutants. Further, expression of *UAS-λhtl* in glia does not affect glial number either. This suggests that in the embryo, the role of Htl in the glia is primarily associated with morphogenesis and not proliferation or differentiation. However, in the larva, expression of *UAS-λhtl* leads to glial proliferation (Avet-Rochex et al., 2012) and mRNA levels of *fog* and *smog* appear to be altered as well (Avet-Rochex et al., 2014), not only supporting our findings regarding an interaction between the Fog and Htl signaling, but also suggesting a context dependent mode of regulation.

It is curious that even though Htl functions as a negative regulator of Fog signaling, the mutants do not exhibit any of the strong CNS phenotypes associated with Fog overexpression. The only resemblance to it is seen in the clustering of glia which indicates increased adhesion – a phenotype also seen in embryos expressing *UAS-fog*. This suggests that Fog signaling is likely to be tightly regulated and Htl is probably one of the many regulators of this pathway.

The interaction between Fog and Htl signaling pathways appears to be through Smog-a GPCR known to bind Fog (Kerridge et al., 2016). Our results show that *smog* functions as a negative regulator of Fog signaling in the CNS. In addition, loss of one copy each of *htl* and *smog* leads to a synergistic enhancement of Fog signaling clearly indicating that they are part of the same regulatory pathway (Fig. 6).

How does one reconcile the interaction between Htl and Smog with the role of Smog as a negative regulator of Fog signaling in the CNS, and its function as a Fog receptor during gastrulation? The answer to this conundrum may lie in the fact that *smog* encodes multiple splice variants which, except for differences in the C-terminal region, are otherwise have identical (Fig. S6). It is possible that the CNS expresses at least two isoforms of Smog which form homo and heterodimers. The active dimer binds and activates Fog signaling while the inactive dimer binds and sequesters Fog thus blocking signaling. We hypothesize that Htl regulates the dynamic equilibrium between these two dimeric states by regulating the levels of one of the isoforms (Smog-Inhibitory/Smog-I) that gives rise to the inactive state. That GPCRs function as dimers is established (Bai, M., 2004; Milligan., 2004) and a recent study by Jha and colleagues has shown that Fog triggers oligomerization of Smog (Jha et al., 2017). It is possible that loss of Htl affects the homeostasis of Smog-I leading to a downregulation of this isoform, which effectively increases not only the number of ‘active receptors’ but also the amount of available ligand, resulting in enhanced signaling. In contrast, constitutive activation of Htl, might function by stabilizing the transducing receptor complex that enables prolonged signaling. The mode for such a regulation in both cases could be phosphorylation.

Phosphorylation by GPCR receptor kinases (Gprks) and other kinases followed by recruitment of beta-arrestins is one of the keys steps in GPCR inactivation and endocytosis (Dewire et al., 2007; Gurevich and Gurevich., 2019). Recent studies on rhodopsin show that phosphorylation patterns on GPCRs can be grouped according to their ability to bind and stabilize, activate, modulate and even inhibit beta-arrestin binding (Mayer et al., 2019). It is therefore conceivable that Htl regulates Smog stability through such a mechanism. Consistent with this, an analysis of the predicted phosphorylation sites in the different Smog isoforms shows presence of a unique pattern in each variant (Fig. S7).

In the model described above, the regulation by Htl would necessarily be context-dependent, based on the identities and expression level of the *smog* isoforms. The fact that variants Smog B, D and F are differentially expressed in the embryo (Fig. S6) lends support to this possibility. It has been difficult to dissect the expression pattern of individual isoforms in the embryo using in-situs given the strong nucleotide sequence identity amongst the splice variants. *SmogC* which encodes one of the longer isoforms of *smog*, has been shown to bind and mediate the Fog signal during gastrulation (Kerridge et al., 2016). Supporting this, expression of *smogC::GFP* in the CNS enhances AMC due to Fog overexpression (Fig. S3). Based on sequence comparison of all the isoforms, we predict that SmogB is likely to function as a potential negative regulator given its very short C-terminal domain. However, this will need to be tested.

Our results thus support a role for FGFR/Htl in modulating and fine-tuning Fog signaling in a threshold dependent manner. At a low signaling threshold, Htl could potentiate Fog signaling by affecting the stability of the negative regulator whereas at a high threshold, it could enhance signaling by stabilizing the signaling complex. Whether this is indeed so, will need to be tested through future studies.

## Materials and Methods

### *Drosophila* stocks and fly husbandry

All fly stocks were raised on standard cornmeal medium. *htl^AB42^/TM3[ftz::lacZ] (*#5370), *UAS-λhtl* (#5367), *UAS*-*ctaRNAi* (#51848), *UAS*-*htlRNAi (*#35024), *C155*-GAL4 (#458), *UAS-CD4-td-GFP* (#35839), *UAS-mistRNAi* (#41930), *UAS-htl (#5419), smog^MI04401^(#37662)* are from the Bloomington *Drosophlla* Stock Centre, USA. *UAS*-*smog* RNAi (GD7852) is from the VDRC stock center, Vienna, Austria. *htl^YY262^/TM3[ftz::lacZ] (Gisselbrecht et al., 1996); dof^111^* (M.Leptin, Univ.of Cologne, Germany); *UAS-cta, UAS-cta^Q303L^ and cta^RC10^* (Naoyuki Fuse, NIG, Japan); *Smog^KO^* and *UAS-SmogC::GFP* (S.Kerridge and T.Lecuit, AMU, France); *htl* ^s1-28^/TM3 (T.Kojima, Japan); *htl*-GAL4 (Alicia Hidalgo, Univ.of Birmingham, UK); *UAS-ths::HA* (Arno Muller, UK); *UAS-ths* (Angelike Stathapolous, Caltech, USA); *elav*-GAL4 and *W^1118^* (K. Zinn, Caltech, USA); *C155*-GAL4; *elav*-GAL4, *UAS-fog* and *UAS-fogRNAi* used in this study have been described previously (Ratnaparkhi and Zinn, 2007). Except where stated, all experiments were carried out at 25°C. Balancers carrying lacZ or GFP were used to identify animals of the correct genotype. For all crosses involving *C155*-GAL4; *elav*-GAL4, the control was *C155*-GAL4/+; *elav*-GAL4/+. This has been referred to in short as ‘*C155*-GAL4; *elav*-GAL4/+’ in the text. The genotypes of all the lines generated and used in this study have been included as supplementary material. For the experiment involving *UAS-mistRNAi*, we recombined *UAS-fog* and *UAS-mistRNAi*. We validated the line for presence of *UAS-mistRNAi* by crossing the recombinant line to *C155*-GAL4; *elav*-GAL4, and checking for downregulation of Mist in larval brains using anti-Mist. We confirmed the presence of *UAS-fog* by staining *elav-GAL4>UAS-fog, UAS-mistRNAi* embryos with anti-Fog.

### Generation of Smog antibody

The region corresponding to the first 78 residues in the N-terminal region of Smog was amplified by PCR using primers containing EcoRI and XhoI site in the forward and reverse primers respectively. Forward primer: 5’ AG GAA TTC ATG GAA CTG TGC ATA GC 3’; Reverse Primer: 5’ GA CTC GAG GTT GCA CCT AGT GGA T 3’ The PCR product was first cloned into pGEM-T easy vector (Promega), verified by sequencing and subsequently cloned into pGEX5.1 vector using the EcoRI and XhoI sites. Purified GST tagged protein was used to raise antibodies in rabbits (Chromous Biotech, Bangalore, India). We have tested the antibody using different dilutions and found that it works well in a broad range of dilutions. We have tested pre-adsorbed antibody at 1:600 and unadsorbed antiserum at 1:2000. Under both conditions, we have obtained similar results in our experiments without any change in the staining pattern. For most of our studies here we have used the pre-adsorbed antibody at a dilution of 1:600.

### Immunohistochemistry and Image Analysis

Embryo fixation and immunohistochemistry were performed using standard protocols (Patel, 1994). Embryos of the correct genotype were scored using lacZ or GFP balancers. The following antibodies were used: chicken anti-GFP (1:1000, Life Technologies), rabbit anti-GFP(1:1000, Life Technologies), mouse anti-GFP (1:1000, Life Technologies), rabbit anti**-**Beta Galactosidase (1:1000, Life Technologies), mouse anti-Beta Galactosidase (1:1000, Promega), mouse anti-Beta Galactosidase (1:10, DSHB, Iowa), anti-fasciclin II or mAb1D4 (1:30 or 1:50 DSHB, Iowa), anti-Repo (1:10 or 1:20 DSHB, Iowa), anti-Futsch (22C10, 1:100, DSHB, Iowa), anti-Fog (1:500, N.Fuse, Japan) Secondary antibodies conjugated to Alexa Fluor 488, 568, or 633 (Molecular Probes) were used at 1:1000 (1:1000, Life Technologies). For all immunostainings a common cocktail containing the primary antibodies was made, mixed thoroughly and divided equally between control and experimental tubes. A similar procedure was followed during the addition of the secondary antibody. For the images and quantitation shown in Fig.7, we stained control and mutant embryos in the same tube. Embryos from *htl* mutants and *w^1118^* embryos were first individually stained for the balancer using anti-Beta galactosidase or anti-GFP antibodies. Mutant embryos at stage 16 or late stage 16 were sorted on the basis of balancer staining. An approximately equal number of wild type embryos of the same stage also were sorted. During sorting, the anterior region of one and the posterior region of the other, was cut to mark the genotype; the embryos were mixed and stained in the same tube.

Images were obtained using a Leica SP8 confocal system. All glial images were obtained using a 63X objective (N.A = 1.4); a 40X objective (N.A = 1.4) was used to for all axonal images. Abdominal segments A1-A7 and A2-A7 were used to quantify axonal midline crossing and glial midline crossing respectively. While imaging for measuring intensity, we have used identical confocal settings for control and experimental samples.

### Geometric Measurements

Aspect ratio was measured using the ImageJ software (NIH). For a given glia, individual ’z’ sections were scanned and the section in which the glia appeared to be of maximum size was selected. The outline of the cell was drawn manually using Repo staining (nuclear) and the membrane-GFP staining as markers. The ‘fit ellipse’ and ‘shape descriptor’ tool in imageJ was used to obtain values for aspect ratio. For each embryo 20 dorso-medial LG were randomly selected for analysis; 20-25 embryos were used for each genotype. The values obtained for aspect ratio were normalized to their respective control. These indices are unitless.

### Measurement of Smog intensity

The intensity of Smog staining at the chordotonal organ was quantified using the Image J software (NIH, Bethesda). We have measured the intensity of Smog staining for each individual neuron by manually selecting a single confocal section that showed the highest staining intensity for Smog for a given neuron. The stained area at the tip of the dendrite was manually outlined and the mean grey value for intensity was measured using the Image J software. The intensity in the nuclear region was taken as a measure of the background staining since we tend to see significant Smog staining in the muscles and cells surrounding the chordotonal organ. The average background value was calculated from the intensity of at least 5 distinct nuclei from independent chordotonal organs in a given embryo. This value was subtracted from the Smog intensity obtained for individual neurons for that embryo. The mean gray value after background subtraction was normalized and used to plot the graph.

### Quantitative PCR

Stage 16 and stage late 16 embryos were used to isolate total RNA using the Trizol method according to the manufacturer’s instructions. Homozygous *htl^AB42^* mutants were sorted unequivocally based on the absence of the GFP balancer and also on the basis of the gut defects seen in these mutants. cDNA synthesis was carried out from 1μg of total RNA using Verso cDNA synthesis kit from thermoscientific (Cat. No.AB-1453A). Quantitative PCR was carried out using QuantStudio3 (Applied Biosystems) using SYBR Green GoTaq master mix from Promega (Cat.No. A6001). Calculation of relative gene expression was done after normalization to control *rp49* using the *ΔΔ*Ct analysis method. The graph shown is the average obtained from 3 biological replicate. The value for each biological sample was an average of 3 technical replicates. The sequence of the primers used are as follows:

*smogB* Forward: 5’-GTTTCGCCTTGGGTCTGATA 3’

*smogB* Reverse: 5’ GGTCTGTGCTTATTGGTCGT 3’

*smogD* Forward: 5’ GCAGAAAAAGGGCAGCAA 3’

*smogD* Reverse: 5’ CCTCGGTGATCTCGATGT 3’

*smogF* Forward: 5’ AACCTTGAAAGCAGCCAAGA 3’

*smogF* Reverse: 5’ ATCGAACTCAAGACTGACAG 3’

*htl* Forward: 5’ CAAGCGGATCGCTGGTAGTG 3’

*htl* Reverse: 5’ GACTCCTGGCTCCCAAATGAT 3’

### Statistics

Statistical analyses was done using the GraphPad Prism Software. All comparisons were made using student’s *t*-test. The scatter dot plots are represented as mean±SD. All other bar graphs are mean± SEM.

## Acknowledgements

We thank BDSC and VDRC stock centers for stocks; DSHB, Iowa State for antibodies; Steve Kerridge, Maria Leptin, Naoyuki Fuse and Steve Rogers for fly stocks and antibodies; Bhagyashree Kaduskar and Kavita Babu for critical comments; KS was supported by a fellowship from the University Grants Commission, Govt. of. India; AB was supported by a senior research fellowship from CSIR, Govt. of India. We gratefully acknowledge funding support from SERB, Govt. of. India and intra-mural funds from MACS-ARI (ARI/ZOO16) to AR.

## Figure legends

**Fig. S1.**
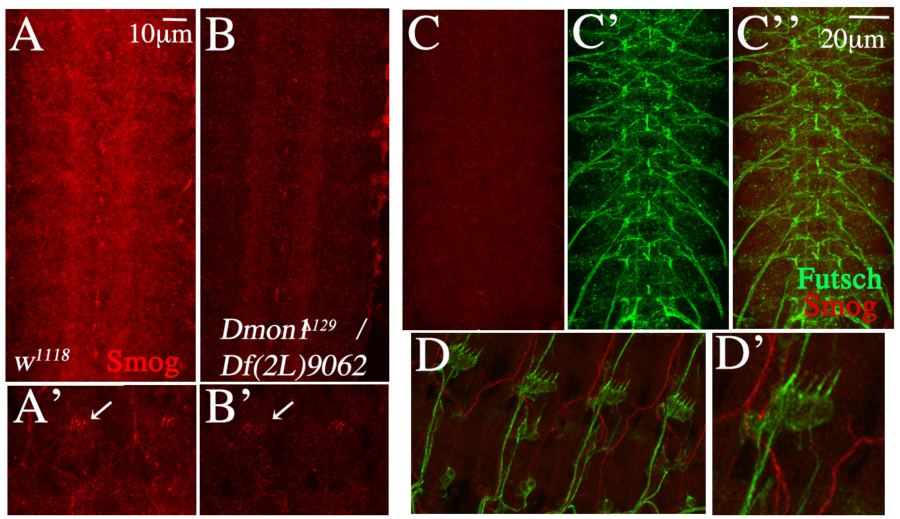
Analysis of Smog staining in the embryonic CNS. (A-B) CNS and PNS of wildtype (A, A’) and *Dmon1^Δ129^/Df(2L)*9062 (B, B’) embryo stained with the anti-Smog (red). A decrease in staining intensity is observed in the mutants. (C-D) Wild type embryo stained with pre-immune serum (red) and anti-Futsch (22C10, green). No staining is observed in the CNS (C-C’’) and in the chordotonal organ (D-D’).

**Fig. S2.**
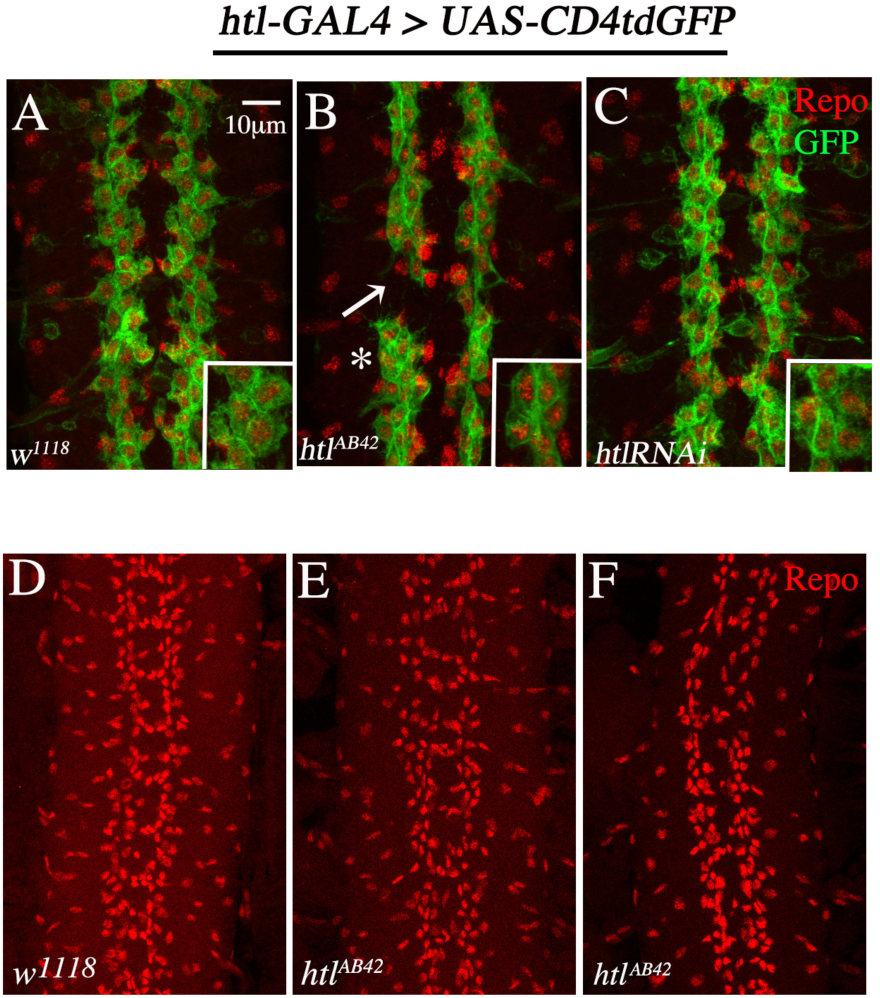
Glial morphology and organization in *htl* mutants. (A) CNS of a control embryo (*htl*-GAL4, *UAS-CD4tdGFP/+*) stained with anti-Repo (red) and anti-GFP (green) antibodies. LG are oval and flat (inset). (B) LG in *htl*^AB42^ mutants appear clustered (asterisk, see inset) and show occasional gaps (arrow) between two segments. (C) LG in embryos expressing *htlRNAi*, tend to cluster and appear smaller in size. (D) LG in wildtype embryos at late stage 16 appears organized in two longitudinal columns. (E-F) LG show mild disorganization in *htl^AB42^* mutants.

**Fig. S3.**
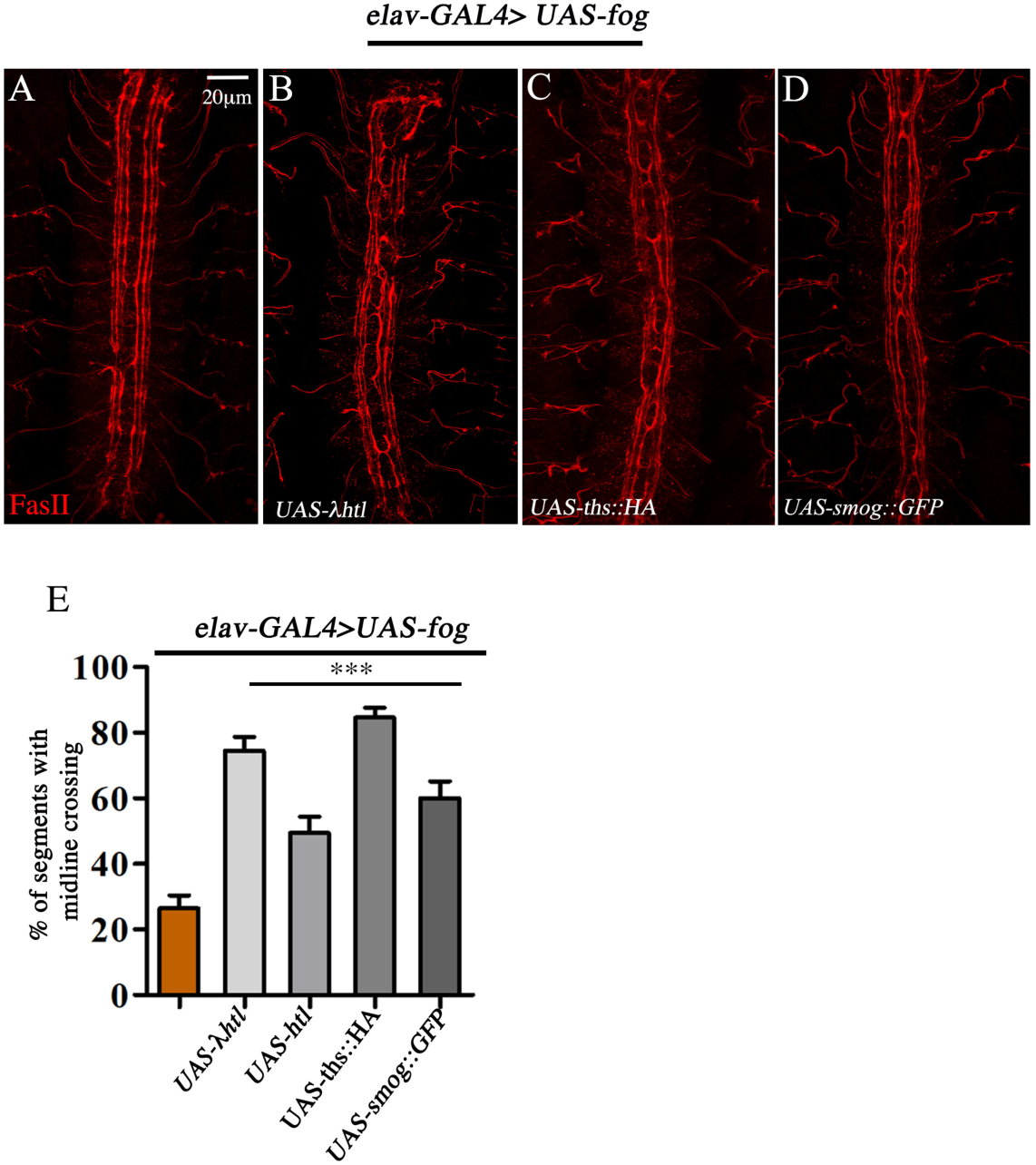
Constitutive activation of Htl signaling enhances AMC. (A) Expression of *UAS*-*fog* in neurons leads to ectopic axonal midline-crossing (AMC). (B-C) Co-expression of either *UAS-λhtl* (B) or *UAS-ths::HA* (C) with *UAS-fog* enhances AMC. (D) Expression of *UAS-SmogC::GFP* also enhances midline crossing. (E) Quantification of AMC: *elav*-GAL4>*UAS-fog* (26.4±4.02, n=182); *elav*-GAL4>*UAS-fog;UAS-λhtl* (74.53±4.21,n=161); *elav*-GAL4>*UAS-fog;UAS-htl* (49.4±5.0,n=168); *elav*-GAL4>*UAS-fog;UAS-ths::HA (84.62±2.96, n=182); elav*-GAL4>*UAS-fog;UAS-SmogC::GFP* (60.0±5.13, n=70). All values are mean±SEM; n=number of segments. P<0.0001

**Fig. S4.**
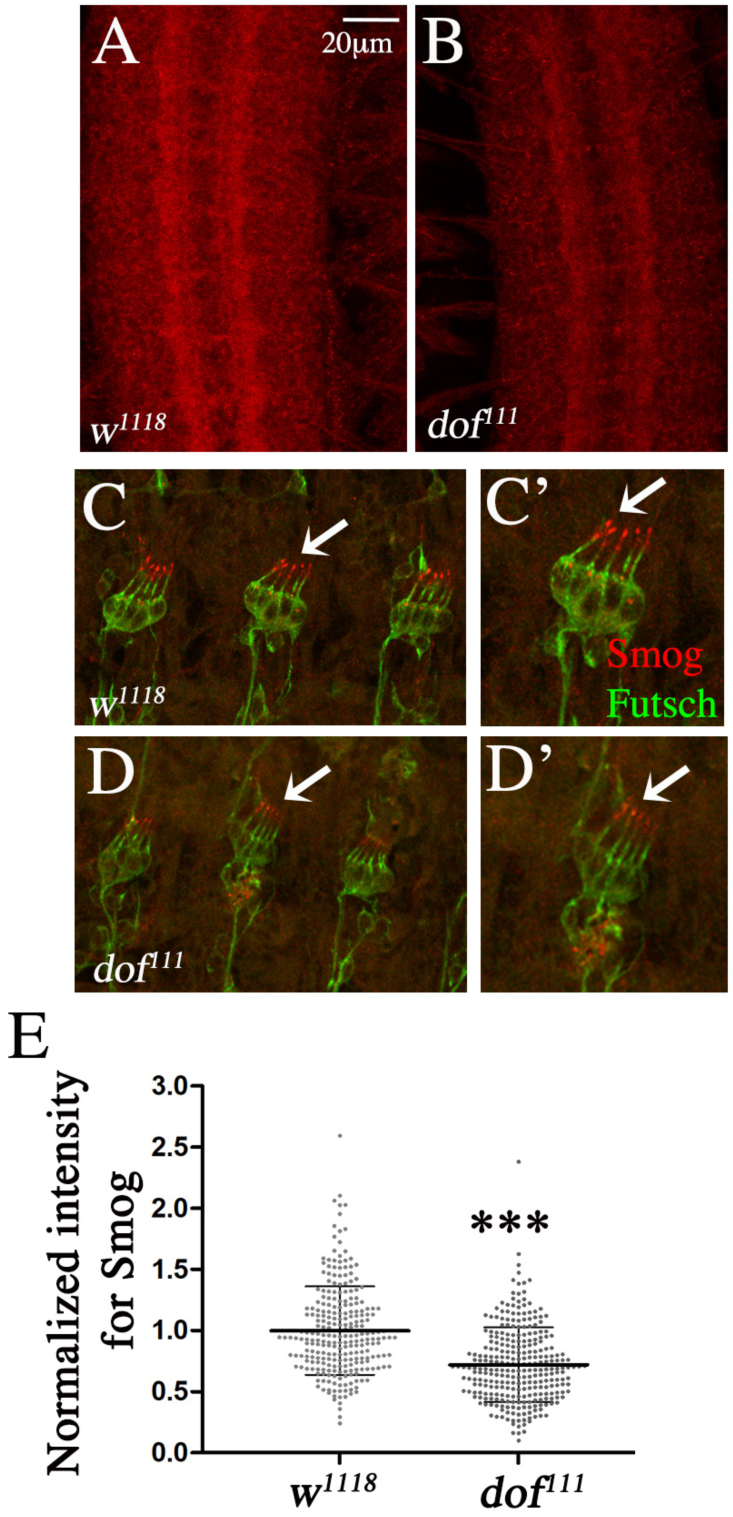
Smog staining is reduced in *dof* mutants. (A&B) CNS of wildtype (A) and *dof^111^* mutant (B) stained with anti-Smog. Note the decrease in Smog staining in the mutant. (C-D) Smog staining in the chordotonal organs is reduced in *dof^111^* mutants (compare C, C’ to D, D’). (E) Graph of the staining intensity in the chordotonal organs shows a 28% decrease in *dof^111^* mutants (w^1118^: 1 ± 0.36, N=17, n=260) versus *dof^111^*: 0.72±0.31, N=17, n=291) N= number of embryos; n= number of neurons; *** indicates P<0.0001.

**Fig. S5.**
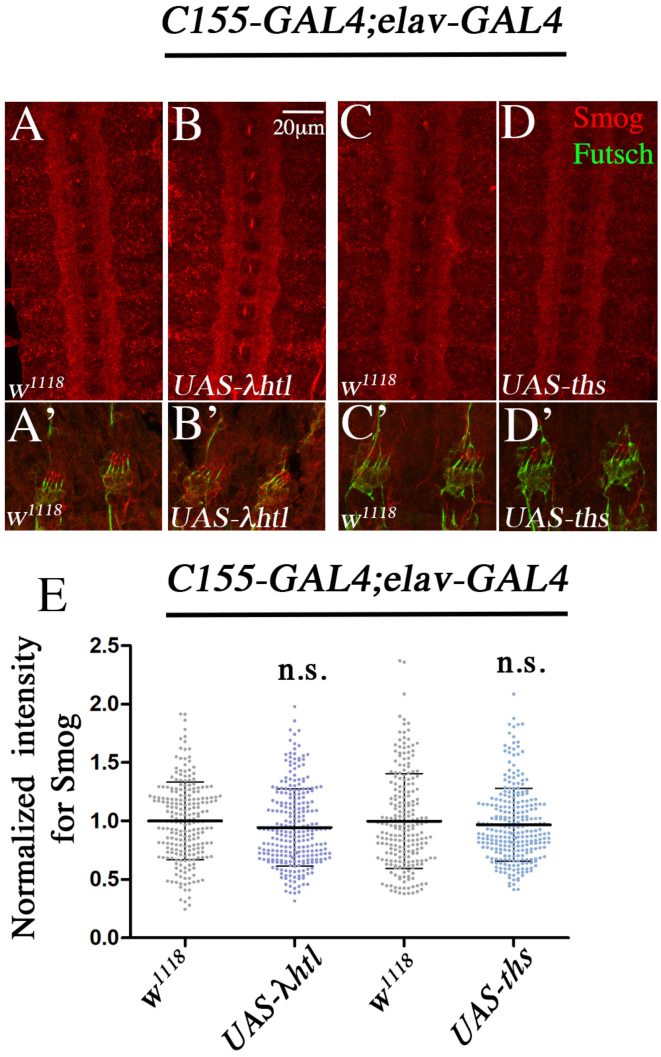
Ectopic activation of Htl signaling does not affect Smog levels. Shown is the CNS (A-D) and chordotonal organs (A’ –D’) stained with anti-Smog (red, A-D). The green in panels A’-D’ represents staining with anti-Futsch. The intensity of Smog staining does not change upon overexpression of *UAS-λhtl* (Compare A&B and A’&B’) and *UAS-ths* (Compare C&D and C’ and D’). (E) Graph showing normalized mean intensity of Smog staining: *C155*-GAL4; *elav*-GAL4/+ (1.0±0.33, N=12, n=227) versus *C155*-GAL4;*elav*-GAL4>*UAS-λhtl* (0.95±0.33, N=13, n=265); *C155*-GAL4; *elav*-GAL4/+ (1±0.40, N=14, n= 224) versus *C155*-GAL4;*elav*-GAL4>*UAS-ths (*0.97±0.31, N=17, n= 271). Students t-Test and the Mann-Whitney Test was used to analyze the data of *UAS-λhtl* and *UAS-ths* respectively. N= number of embryos; ‘n’= number of neurons.

**Fig. S6.**
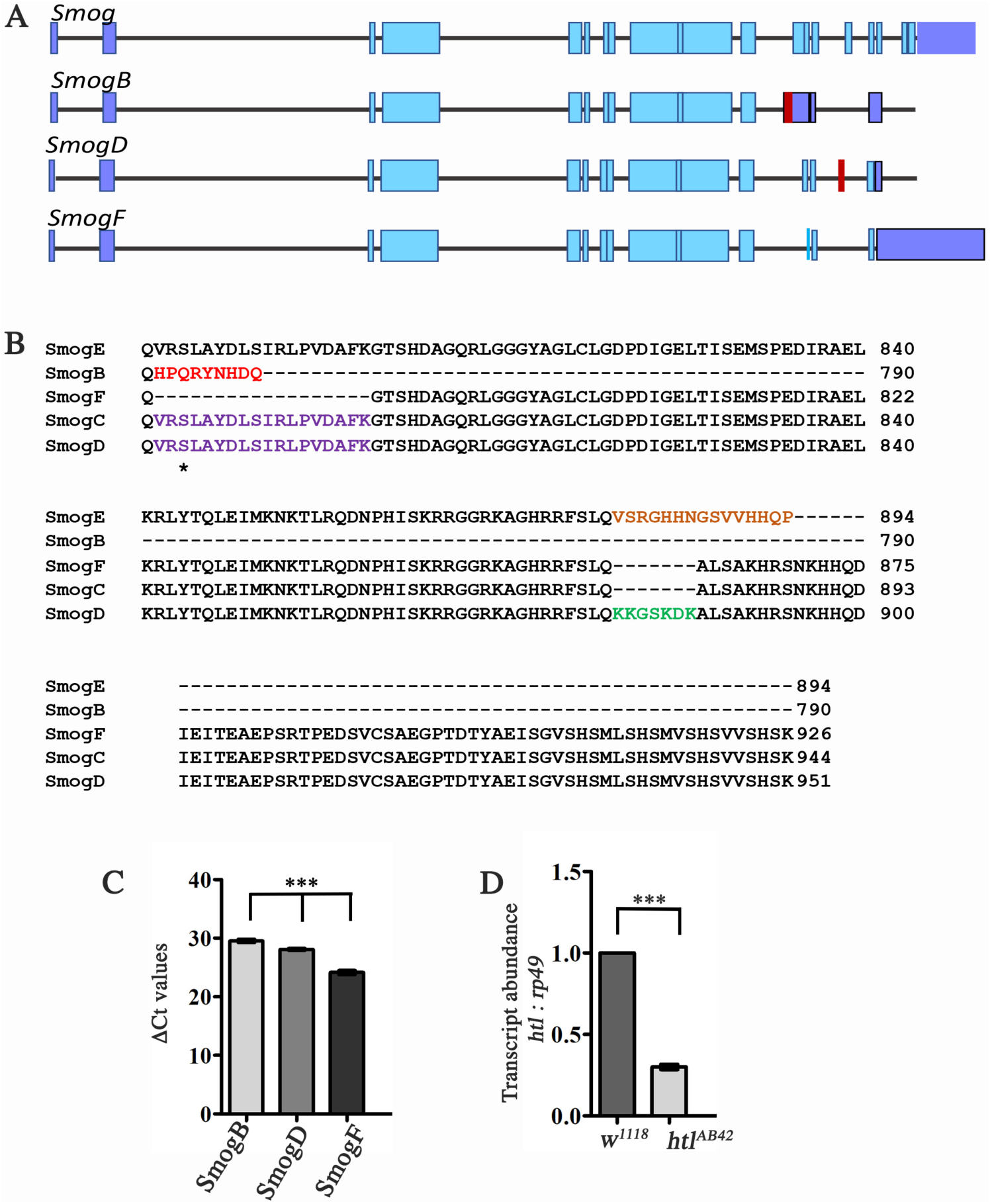
*Smog* isoforms show differential expression in the embryo. (A) Schematic showing the intron-exon pattern of *smog* variants. Unique regions in *smogB* and *smogD* are marked in red. Purple regions refer to the UTRs. (B) Alignment of the C-terminal domain of SmogB, SmogD and SmogF is shown. Residues unique to each variant are marked in color. (C) Shown are the Ct values for *smogB*, *smogD* and *smogF* obtained through qPCR. The value is highest for *smogB* indicating that it is the least abundant of the three isoforms (*smogB*:29.53±0.27; *smogD*: 28.08±0.16; *smogF*: 24.14±0.35). (D) Quantification of the *htl* transcript in *w^1118^* and *htl^AB42^* embryos. The mutants show a 70% reduction in *htl* mRNA (*w^1118^*: 1.0; *htl^AB42^*: 0.30±0.02).

**Fig. S7.**
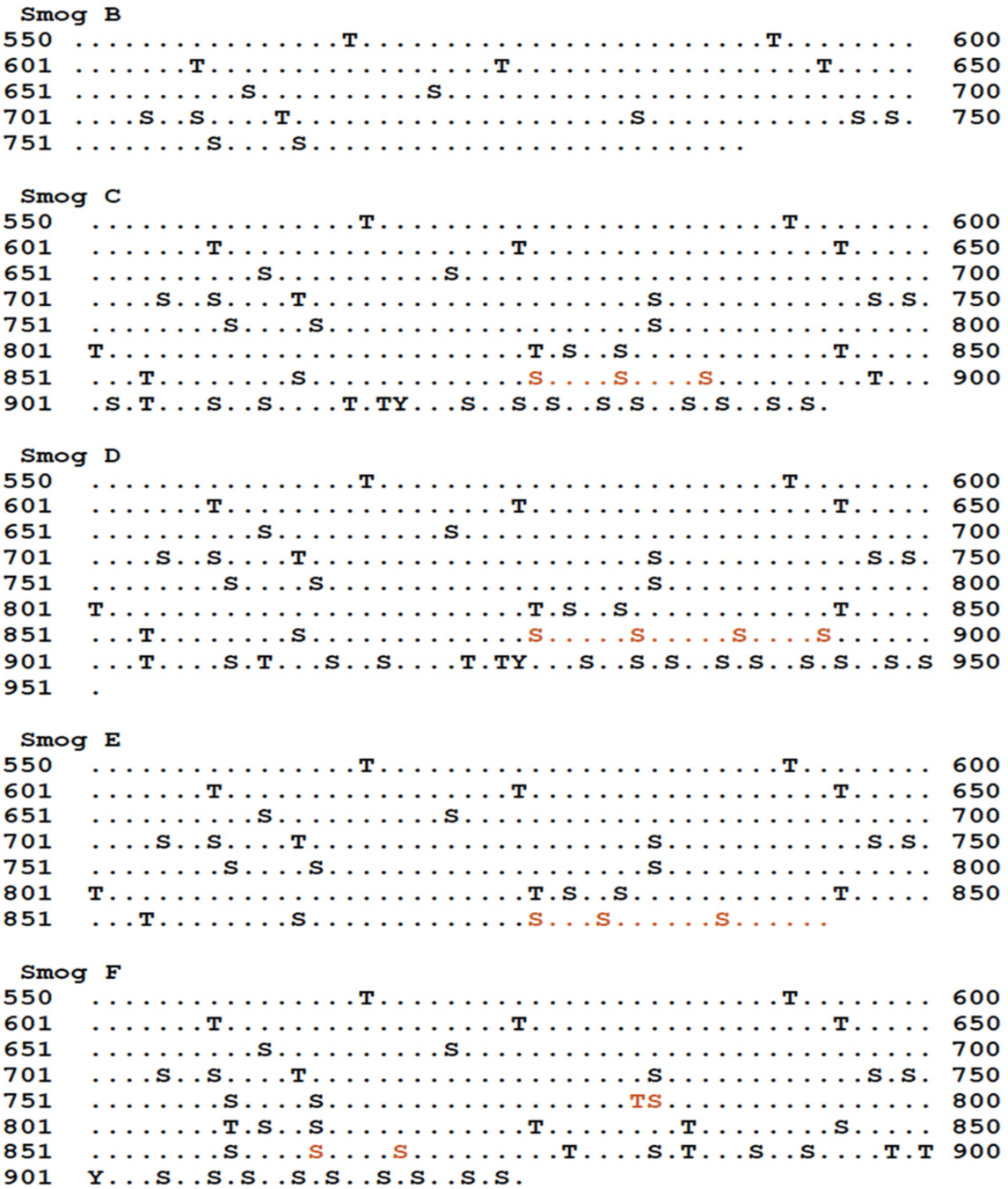
Predicted phosphorylation pattern of Smog isoforms. Shown are the predicted phosphorylation sites for all Smog isoforms. The residues in red are unique to the respective isoform. Thus each isoform could potentially have a distinct phosphorylation pattern that controls its regulation and stability.

